# The preferential injury of outer renal medulla after ischemia-reperfusion relies on high oxidative metabolism

**DOI:** 10.1101/2024.09.12.612245

**Authors:** Grégoire Arnoux, Justine Serre, Thomas Verissimo, Matthieu Tihy, Sandrine Placier, Charles Verney, Frédéric Sangla, Deborah Paolucci, Marylise Fernandez, Sophie de Seigneux, Sebastian Sgardello, Maarten Naesens, Juliette Hadchouel, Éric Feraille, Stellor Nlandu Khodo, Pierre Galichon, David Legouis

## Abstract

Acute kidney injury (AKI) is a prevalent and significant complication in critically ill patients, and its management remains a considerable challenge. The kidney is a highly metabolic organ, consuming and producing substantial amounts of ATP, mainly through mitochondrial oxidative phosphorylation. Recently, mitochondrial dysfunction has been identified as a key factor in the pathophysiology of AKI and the progression to chronic kidney disease. The kidney is a complex organ, comprising millions of structural and functional units. These nephrons are composed of different cell types dwelling within specific metabolic microenvironment. Whether the metabolic spatialization in the kidney has consequences on tubular injury distribution and severity remains unclear.

In this study, we identified the high metabolic rate of the outer stripe of the outer medulla (OSOM) and its substrate preference flexibility, relying on both glycolysis and fatty acid oxidation (FAO) to fulfill its ATP demands. We demonstrated that the OSOM is susceptible to mitochondrial and FAO impairment induced by propofol, the most used sedative in intensive care settings, which exacerbates tubular injury during AKI. In the clinical setting, the cumulative dose of propofol is positively correlated with oxidative metabolism disruption and histological and function outcomes in renal allograft recipients. Finally, we found that the loop of Henle, the OSOM major constituent, was the most injured segment during AKI in patients.

This study shows how renal metabolic spatialization impacts tubular injury severity. We identified the OSOM as the most metabolically active and the most injured region of the kidney both in humans and mice. We demonstrated that propofol is a potent inhibitor of renal mitochondrial function and FAO exacerbating tubular injury in the OSOM upon IRI.

**Translational Statement:**

- Aerobic metabolism is basally enhanced in the renal OSOM, including the S3 proximal tubule, the thick ascending limb of the loop of Henle and Distal Convoluted Tubule
- PT cells as well as TAL cells are significantly targeted by injury in human AKI.
- Propofol impairs renal mitochondrial function worsening tubular injury during ischemia reperfusion.

## Introduction

Acute kidney injury (AKI) is the most common organ failure observed in critically ill patients. The growing incidence of AKI, as reported in the FINNAKI and AKI-EPI studies, reached 39 and 53% respectively, with a quarter requiring renal replacement therapy (RRT)^1–3^. AKI is associated with poor outcomes, including prolonged intensive care unit (ICU) and hospital length of stay, development of chronic kidney disease (CKD), short- and long-term mortality^3–6^ and cardiovascular events^7^. However, the pathophysiology of this devastating syndrome is far from fully understood.

One of the main functions of the kidney is blood purification. For this purpose, the kidney combines two processes: filtration of total plasma and exchange of solutes and water between the filtered blood and the filtrate by secretion and reabsorption. Whilst filtration is driven by blood pressure, reabsorption involves an electrochemical gradient along the nephron generated by active ion pumping. The intact mammalian kidney reabsorbs almost 80 mmol of sodium per day and per gram of kidney^8^. The kidney is thus a highly metabolic organ, containing the largest concentration of mitochondria after the heart, and is characterized by the production and consumption of large amounts of ATP ^8,9^. Under physiological conditions, ATP production mostly derives from oxidative phosphorylation ^10–12^. Recently, metabolic dysfunction has emerged as a key player in the pathophysiology of AKI. In particular, mitochondrial dysfunction and decreased fatty acid oxidation have been shown to promote renal injury and progression to CKD ^13–21^.

Kidneys are sophisticated organs, with each human kidney containing about a million structural and functional units, the nephrons, each of them composed of different cell types. Each cell type tunes its substrate preference and metabolic status depending on its anatomical localization and the energy required to fulfill its function^22^. Energy metabolism is therefore not uniformly distributed along the nephron, with active transport, ATP production and consumption occurring in a coordinated manner^23–28^.

In this study, we investigated the spatial distribution of renal energy metabolism using spatial and single cell transcriptomics and immunofluorescence. Having identified the outer stripe of the outer medulla (OSOM) as the most metabolically active area of the intact kidney, we investigated whether this zone might be vulnerable to inhibition of mitochondrial function during AKI. Finally, we aimed to validate our findings in the clinical setting.

## Methods

### Spatial transcriptomics

Kidneys were harvested from two healthy uninjured C57Bl/6JRj mice and then snap-frozen in liquid nitrogen. Thin sections were cut and transferred to the capture area of the 10X Genomics gene expression slide. After H&E staining and image acquisition, the tissues were permeabilized; reverse transcription and library construction were performed according to the instructions provided by 10X Genomics. The fastq files were processed and aligned on the mouse reference genome GRCm39 using SpaceRanger. The count matrix was further normalized and scaled using Seurat and the SCT method. Within the cortex, the outer and inner strip of the outer medulla (OSOM and ISOM respectively) and inner medulla were delineated using anatomical structures (see annotation of kidney compartment paragraph) and annotated with the Loupe Browser (10X Genomics).

To annotate each spot in terms of cell types, we integrated our spatial transcriptomics dataset with a publicly available renal scRNA-seq dataset^29^, downloaded from Gene Expression Omnibus under accession number GSE139107. To construct our reference atlas, we firstly preprocessed the data in accordance with the conventional Seurat pipeline. Briefly, the data were divided into individual samples. DoubletFinder ^30^ was employed to identify and remove potential doublets. The raw count matrices were individually normalized and scaled using the SCT method from Seurat, and then integrated together using the rpca method. Finally, the runPCA, runUMAP, findNeighbors and findClusters commands were executed. The clusters were annotated using classical gene markers. We further applied robust cell decomposition^31^ to assign cell types to our spatial transcriptomics dataset using the annotated snRNAseq dataset described above. Each spot was therefore annotated as the three most probable cell types predicted.

To assess the activity of the tricarboxylic acid cycle (TCA), fatty acid oxidation (FAO), mitochondrial biogenesis, electron transport chain (ETC), ATP synthesis, gluconeogenesis and glycolysis, we used genesets from the REACTOME database as an input to the singscore algorithm^32^. The fluxes of glucose to lactate and acetyl-CoA to oxaloacetate were estimated using the single-cell flux estimation analysis (scFEA) algorithm, which is capable of inferring the cell-wise fluxome from single-cell RNA sequencing (scRNA-seq)^33^. We also estimated the relative part of anaerobic and aerobic glycolysis as the ratio between the pyruvate to lactate and pyruvate to acetyl-CoA fluxes estimated by scFEA. The pathway activities across compartments were compared using a Wilcoxon test.

### Seahorse Experiments

The Seahorse XF-96 Extracellular Flux Analyzer (Agilent) was employed to quantify oxygen consumption (OCR), proton efflux rate (PER), ATP synthesis rate and substrates oxidation. Renal primary cells were obtained from 6 C57/Bl6 mice after mechanical dissociation performed using the gentleMACS dissociator (Miltenyi Biotec). About 80,000 cells were therefore seeded just after isolation into XF96 cell culture microplates (Agilent) previously coated with CellTak (Corning), with Seahorse XF DMEM (pH 7.4) supplemented with glucose (5 mM), pyruvate (1 mM), and glutamine (2 mM), added or not with propofol (100 µM), and incubated at 37°C in a carbon dioxide-free incubator for 2 hours. Thirty minutes prior to the experiment, the sensor was placed into the XF-96 instrument and calibration was initiated with XF calibrant medium (Agilent).

Baseline OCR and PER were measured after 15 minutes of equilibration. Maximal OCR was calculated as the difference between maximal OCR recorded after FCCP (2µM) injection and minimal OCR recorded after rotenone/antimycinA (0.5µM) injection.

Mitochondrial and glycolytic ATP production was measured using the Seahorse ATP Real-Time rate assay (Agilent) with oligomycin and rotenone/antimycinA used at 3µM and 0.5 µM, respectively.

Fatty acid and pyruvate oxidation was measured using the Seahorse XF Glucose/Pyruvate Oxidation Stress Test and the Seahorse XF Long Chain Fatty Acid Oxidation Stress Test kits with oligomycin (3µM), FCCP (1µM), rotenone/antimycinA (0.5µM), etomoxir (4µM) and uk5099 (2µM).

A similar protocol was used with HK2 with parameters mentioned above. Midazolam, the second most used sedative agent, was used as a second control.

### Mitochondrial Membrane Potential

A single-cell suspension of HK2 was incubated for 210 minutes in DMEM, or DMEM supplemented with propofol (at 100 µM and 400 µM) or FCCP (1 µM) as a positive control. Finally, the Mitotracker Red CMXRos (ThermoFisher Scientific) probe was added at 100 nM and incubated for 30 minutes. Subsequently, the samples were washed, centrifuged, and resuspended in DMEM. DAPI was added at a final concentration of 500 µg/mL, and the single-cell suspension was analysed using a Fortessa cytometer (Becton-Dickinson). Cells were selected based on size using FSC/SSC gating. Singlets were further selected using SSC-H gating. Living cells were selected using DAPI. Finally, the mitochondrial potential membrane was measured on the selected cells (singlets alive). The flow cytometry data was analysed using FlowJo v10 (Becton-Dickinson).

### Ex-vivo Renal Perfusion

To study renal metabolism, we used our previously described ex vivo renal perfusion platform^34^. This platform measures renal oxygen consumption and arteriovenous differences in glucose and lactate. In short, healthy wild-type C57/Bl6 mice were anesthetized with an intraperitoneal pentobarbital injection (150 mg/kg, Inresa) and subsequently euthanized. The superior mesenteric artery, celiac artery, abdominal artery, and inferior vena cava were ligatured to isolate the kidney circulation. Two catheters were then inserted into the vein and aorta respectively, corresponding to the output and input flow. The medium was prepared using PrismaSol B0 (Baxter) enriched with potassium chloride (4 mM), glutamine (1 mM), pyruvate (0.1 mM), human hematocrit (10%), Cernevit (200 mg/L, Baxter), Adaven (1.5%, Fresenius), and bovine serum albumin (20 g/L). The medium was oxygenated with a mixture of oxygen and carbon dioxide (95%/5%) at a flow rate of 2 L/min through a membrane oxygenator (Medos Hilite). The oxygen flow rate was adjusted to maintain the oxygen partial pressure within the perfusion solution at approximately 30 kPa. The kidney received perfusion solution infusion from the aorta to the inferior vena cava at a rate of 700 µL/min. Following a 20-minute period of kidney flushing, the perfusion flow rate was decreased to 350 µL/min, and samples were obtained sequentially through the two catheters.

Samples were analyzed using the ABL90 Flex analyzer from Radiometer. Before and after change in renal arteriovenous contents were compared using a linear mixed model with random intercept.

### Animals

Eight-week-old male C57Bl/6JRj mice (weight 25 +/- 2.5 g) purchased from Janvier Lab were used in all experiments. All mice were housed with a 12/12h light-dark cycle with ad libitum access to food and water. All experimental procedures were performed in accordance with the Directive 2010/63/EU of the European Union. Experimental groups were kept in different cages.

### Ischemia–Reperfusion Injury With RIRI Clamping Device

The previously described RIRI clamp method was employed to induce renal ischemia reperfusion injury^35^. On the day prior to ischemia, the mice were anesthetized with an intraperitoneal injection of a solution containing ketamine and xylazine (100 mg/kg and 10 mg/kg, respectively). Following the induction of anesthesia, the left (for the RIRI positioning) and right (for nephrectomy) sides of the abdomen were shaved and disinfected with Betadine® solution. The animal was kept on a heating pad set at 37 °C. For the nephrectomy, the right kidney was exteriorized and excised after ligature of the vascular pedicle. Subsequently, the right kidney was exteriorized, and the renal vascular pedicle was carefully dissected, exposing the blood vessels. The thread of the RIRI clamp was placed around the pedicle with the two cylinders on each side of the pedicle. The kidney was then released into the abdominal cavity. The catheter was inserted into the posterior incision with the flexible end towards the kidney. The extremities of the thread were then inserted within the tube of the catheter, which was exposed through the other end and blocked by the cap of the catheter. Following completion of the surgical procedure, the mouse received a subcutaneous injection of buprenorphine (0.05 mg/kg).

### Sedation and Ischemia

The mice were lightly sedated by exposure to 2% isoflurane on a heating pad the day after clamp-positioning surgery. Once anesthesia had been stabilized at 1% isoflurane, one group of mice received an intraperitoneal injection of propofol (150 mg/kg, n=16), and the control group received an intraperitoneal injection of midazolam (20 mg/kg, n=16). The preconditioning period lasted 60 minutes, during which isoflurane administration was maintained to ensure equivalent depth of anesthesia. Following the 60-minute period, the cap of the catheter was removed, and the clamp was closed by pulling the external extremities of the thread. Once the clamp was in tension, the cap was closed. After 25 minutes of ischemia, the catheter cap was removed to release the tension of the thread, and then closed again. The mouse was maintained in the neonatal incubator at 28 °C until the conclusion of the experimental procedure (the following day) to prevent any decrease in body temperature. With a hypothesis of a 0.05 mL/min difference in GFR between groups, a GFR standard deviation of 0.05 mL/min, a power of 0.8 and an alpha risk of 0.05, the required sample size was 15 animals per group. We enrolled 18 animals per group in 3 different experience batches. In the final analyses, 16 and 14 animals in the propofol and midazolam groups, respectively, were analysed, while 2 and 4 animals in the propofol and midazolam groups, respectively, died before day 2. All animals alive on day 2 were included in the analyses, without exclusion. Simple complete randomisation was used to assign each mouse to the experimental group. During the experiment, mice from the midazolam and propofol groups were alternated to reduce confusion. The people who measured the outcomes (*i.e.* renal function and histology) and performed the analyses were blinded.

### Transcutaneous GFR Measurement

FITC-sinistrin clearance was used to measure GFR as previously described^34^. Briefly, a mini-camera was attached to the mouse flank and a solution of FITC-sinistrin (Fresenius Kabi) was injected into the tail at 0.35 g/kg after 2 minutes of basal recording. The recording of FITC-sinistrin was analyzed using MPD Lab software (Mannheim Pharma and Diagnostics) and GFR was calculated using the appropriate formula^36^.

### Histological Analyses

The kidneys were fixed in formalin 10% for four hours and embedded in paraffin. Sections (3-μm thick) were processed for immunofluorescence. Following deparaffinization and rehydration, antigen retrieval was conducted by incubation for 30 minutes in the target retrieval solution (pH 6 for anti-HK1, anti-ATP5A and anti-megalin; pH 9 for anti-TFAM and anti-megalin (S2369 and S2367, Dako, Agilent Technologies, Santa Clara, CA) at 100°C in a pressure cooker. Subsequently, the sections were incubated in phosphate-buffered saline (PBS) containing 0.1% Triton X-100 and 10% bovine serum albumin (BSA) for 45 minutes. The sections were incubated overnight in a humidified chamber at 4°C with the following antibodies: rabbit anti-TFAM (1:100, PA5-68789, Invitrogen, Waltham, USA), goat anti-megalin (1:2000, gift of Pr. R. Kozyraki^37^), rabbit anti-hexokinase 1-Alexa Fluor 647 (1:100, ab197864, Abcam, Cambridge, UK), mouse anti-ATP5A (1:500,ab14748, Abcam, Cambridge, UK) and rabbit anti-TFAM (1:100, PA5-68789, Invitrogen, Waltham, USA), diluted in PBS-BSA 3%. The following day, after washes in PBS, sections were incubated for one hour at room temperature with Alexa Fluor 594-conjugated donkey anti-rabbit IgG secondary antibody or Alexa Fluor 647-conjugated donkey anti-mouse IgG secondary antibody or Alexa Fluor 488-conjugated chicken anti-goat IgG secondary antibodies, each diluted at 1:500 in PBS-BSA 3%. Subsequently, the nuclei were counterstained with DAPI (1:4000, 62,248, Thermo Fisher Scientific, Waltham, Massachusetts) for 5 minutes. Images were obtained using the slide scanner Axioscan Z1 (Carl Zeiss, Jena, Germany).

### Annotation of kidney compartment

Kidney regions of interest were delineated using anatomical structures (cortex, glomeruli and arcuate vessels; outer stripe outer medulla (OSOM), transition straight portion of the proximal tubule and thin descending limb of Henle; inner stripe outer medulla (ISOM), thin descending/ascending limbs of Henle and collecting ducts; inner medulla, urothelium part of the collecting ducts) in a blinded manner on each section by two experienced nephropathologists (GA and MT).

### Tubular Necrosis Quantification

Digitalized Masson’s trichrome stained sections were further processed and analyzed using QuPath (0.5.1) open-source software^38^ and a script based on a trained pixel classifier was applied. The extent of affected tubular necrosis area was expressed as a percentage of affected areas per region surface of interest (cortex, OSOM, ISOM and inner medulla).

### Immunofluorescent image quantification

Cells were detected using the QuPath’s DAPI algorithm-based cell detection tool, using the optical density sum to segment nuclei. Cells were then classified with the “Train Object Classifier” tool using a neuronal network trained on 20-40 annotations of each cell class (positive or negative cell). Object classifier algorithm was then applied to each slide of the dedicated condition. Results are expressed as a percentage of positive cells per all nuclei of the area of interest.

For intensity quantification experiments, following cell detection, binary threshold-based classification maximal “cellular” megalin positive and negative cells was applied and Atp5a channel cellular max intensity for each cell was obtained.

### Allograft Kidney Recipients

A total of 42 allograft kidney recipients were enrolled at the University Hospitals of Leuven. For each patient, a renal protocol biopsy was performed at four different time points: before implantation (the kidney was flushed and stored in ice), after reperfusion (at the end of the surgical procedure) and 3 and 12 months after transplantation. Genome-wide gene expression profiling using RNA-seq was performed in kidney allograft recipients as previously described^39^. Libraries were prepared using Clontech SMARTer technology and sequenced using the HiSeq 3000 system on the Illumina platform, with a target of 30 million reads per sample. The reads were aligned to the Ensembl top-level assembly with STAR version 2.0.4b. To assess the activity of FAO, ETC, TCA, glycolysis and mitochondrial biogenesis, we employed genesets from the REACTOME database as an input to the singscore algorithm. A robust linear model was used to assess the association between the cumulative dose of propofol and the change in metabolic activity between pre- and post-reperfusion periods. The association between the cumulative dose of propofol and the CT Banff score at one year was evaluated using a Wilcoxon test.

### Kidney Biopsies in AKI Patients

Kidney biopsies, nuclei isolation, snRNAseq and downstream analyses were performed as previously described^40^. Briefly, kidney biopsies were performed just before the therapeutic withdrawal once the consent was obtained from the next of kin. Nuclei were then isolated and snRNAseq was performed. Data were integrated with 3 controls from available datasets generated with similar tissue processing and single-cell technology, downloaded from GEO at the accession number GSE131882. After data integration, major cell types were identified based on gene markers. Detection of differentially abundant cell subpopulations was further performed using DAseq algorithm^41^.

### Statistical analyses

Continuous numerical data were compared using a Mann-Whitney test. All analyses were performed using R software. P-values were two-tailed and a value less than 0.05 was considered significant.

### Ethics Approval

All experimental procedures were performed in accordance with the Directive 2010/63/eu of the European Union. The study received approval from the Ethics Committee Charles Darwin Pitie Salpetriere Group (#27761).

Concerning the cohort of allograft kidney recipients, all patients gave written informed consent, and the study was approved by the Ethical Review Board of the University Hospitals of Leuven (S53364 and S59572).

Concerning the ICU patients, the study was approved by the local ethical committee for human studies of Geneva, Switzerland (CCER 2020-00644, Commission Cantonale d’Ethique de la Recherche) and performed according to the Declaration of Helsinki principles. Consent was given by the next of kin.

## Results

### Aerobic metabolism is basally enhanced in the renal OSOM

To investigate the spatial organisation of renal energy production, we performed spatial transcriptomics on kidneys from two healthy mice. We identified the renal anatomical layers including the cortex, the inner and outer stripe of the outer medulla (OSOM and ISOM) and the inner medulla using a morphological annotation based on histological structures (**Figure 1a**). Concurrently, we conducted an unsupervised clustering analysis on the spatial transcriptomics dataset and observed a complete overlap between the two methods (**Supplemental Figure 1**). We then estimated the score level of the mitochondrial electron transport chain (ETC) based on transcriptomic data. This pathway was expressed at a higher level in the OSOM than in the other regions of the kidney (**Figure 1b-c**). Similarly, mitochondrial biogenesis, and fatty acid oxidation (FAO) were expressed at elevated levels in the OSOM (**Figure 1c**). ATP synthesis by Complex V was also highly expressed in the outer medulla, but at a higher level in the inner strip. As previously described, gluconeogenesis essentially occurred in the cortex, whereas glycolysis, including aerobic and anaerobic glycolysis, was increasing from the cortex to the ISOM. (**Figure 1c**). Interestingly, we compared the relative contribution of anaerobic glycolysis reflected by the estimated glucose-to-lactate flux and the estimated TCA flux and found that TCA was dominant and stable in the cortex and OSOM. In the ISOM, anaerobic and aerobic pathways were equally active. Finally, anaerobic glycolysis was predominant in the inner medulla. (**Figure 1d**). To support these findings, we performed immunofluorescence on renal tissue to stain the ATP synthase F1 subunit alphaprotein, a subunit of mitochondrial complex V. Quantification of the staining intensity level showed higher expression of Atp5a in the outer medulla compared with the cortex, with inner strip displaying a higher expression level than the outer strip (**Figure 1e,f**) following the ATP synthesis activity previously described (**Figure 1c**).

**Figure 1:**
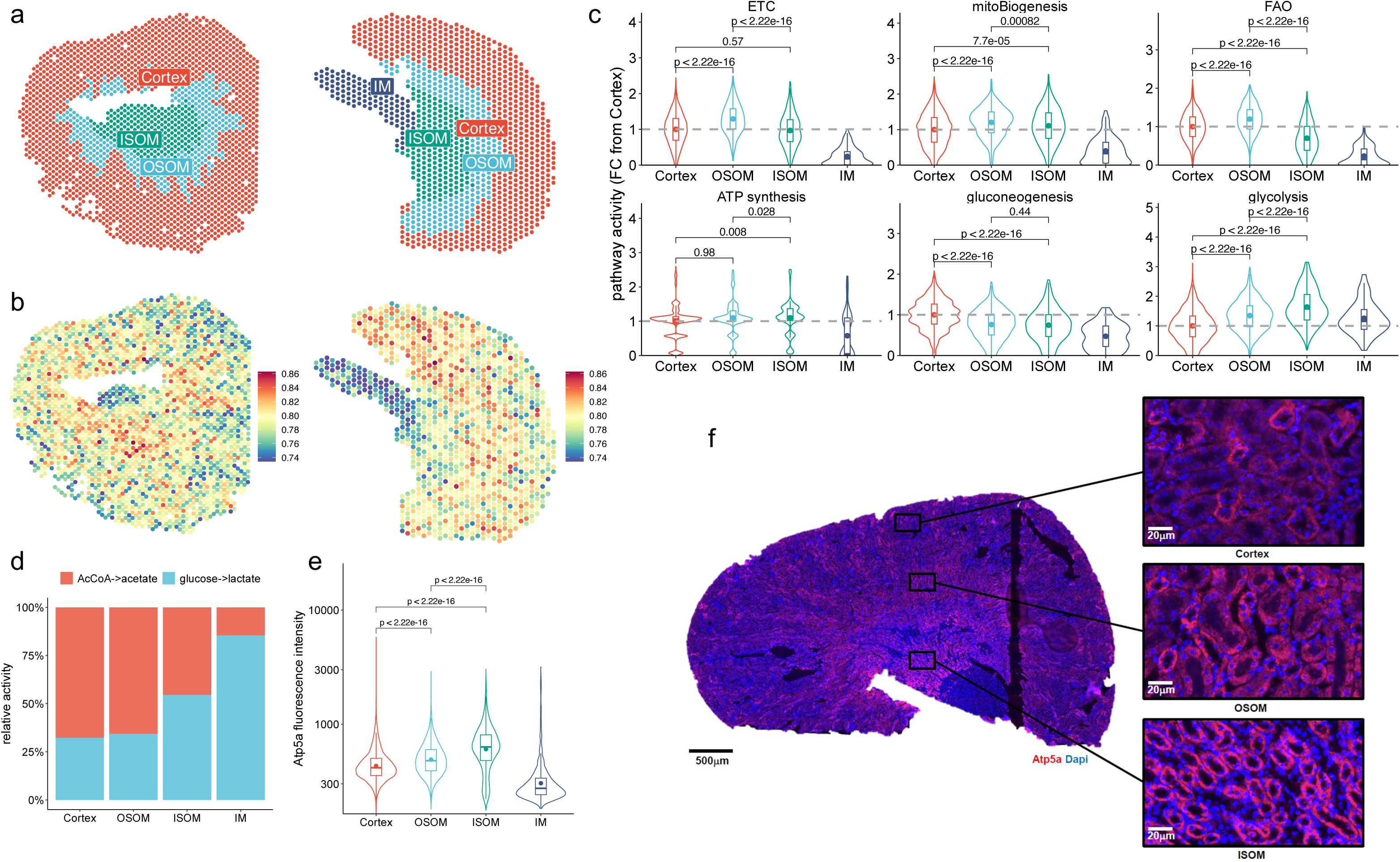
Spatial landscape of the renal energy metabolism. **a**) Spatial transcriptomics morphological annotation depicting the 4 studied compartments. **b**) Spatial transcriptomics pathway activity analysis of the mitochondrial electron transport chain (ETC). **c**) Violin plots showing the activity of different metabolic pathways per compartment in FC from Cortex. **d**) Barplots showing the relative proportion of anaerobic glycolysis and TCA cycle in energy production for each compartment. **e**) Fluorescence intensity level of the Atp5a staining in each compartment **f**) Representative immunofluorescence image of Atp5astaining in healthy mouse kidney section with a focus on cortex (C), outer/inner stripe of outer medulla (respectively OSOM and ISOM). Violin plots display distribution of the data, boxplots display median with interquartile range and dots display mean.

These findings indicate the high metabolic activity of the outer medulla, with the outer strip involving FAO and aerobic glycolysis, while the inner strip relies on aerobic and anaerobic glycolysis.

### Aerobic metabolic processes are basally enhanced in S3 segment, TAL and DCT

We then aimed to identify the cell type with the highest energy production. Therefore, we applied a robust cell type decomposition algorithm. For this purpose, we annotated a publicly available renal scRNAseq dataset ^29^, identifying all major renal cell types based on known gene markers. This dataset was used as a reference to decompose each spot of our spatial transcriptomics dataset (**Figure 2a, Supplemental Figures 2 and 3**). As expected, each renal compartment showed a distinct cell composition, with proximal tubules representing the main cells in the cortex, S3 proximal tubule (PTS3) and thick ascending limb (TAL) in the outer medulla (OSOM and ISOM), and thin limb of the loop of Henle and collecting duct in the inner medulla (**Figure 2b**). Finally, we examined the metabolic pattern at the cell type level. PTS3, distal and connecting tubules (DCT-CNT) and TAL were identified as the most metabolically active cell types (**Figure 2c**), relying on major oxidative processes including aerobic glycolysis and anaerobic glycolysis for ISOM TAL. Anaerobic glycolysis was also high in the collecting duct and the thin limb, two segments part of the inner medulla. To confirm these findings at the protein level, we co-labeled ATP5A and megalin proteins on a kidney section. Megalin, a marker of the proximal tubules, was mainly expressed in the cortex and OSOM (**Figure 2d, e**). Interestingly, the intensity level of ATP5A was higher in the megalin-negative cells in both cortex and OSOM (**Figure 2f**), suggesting that non-proximal tubule cells display a higher mitochondrial content in these compartments. Altogether, these results highlight the high energy expenditure of PTS3, TAL and DCT-CNT, the first two segments being located in the OSOM.

**Figure 2:**
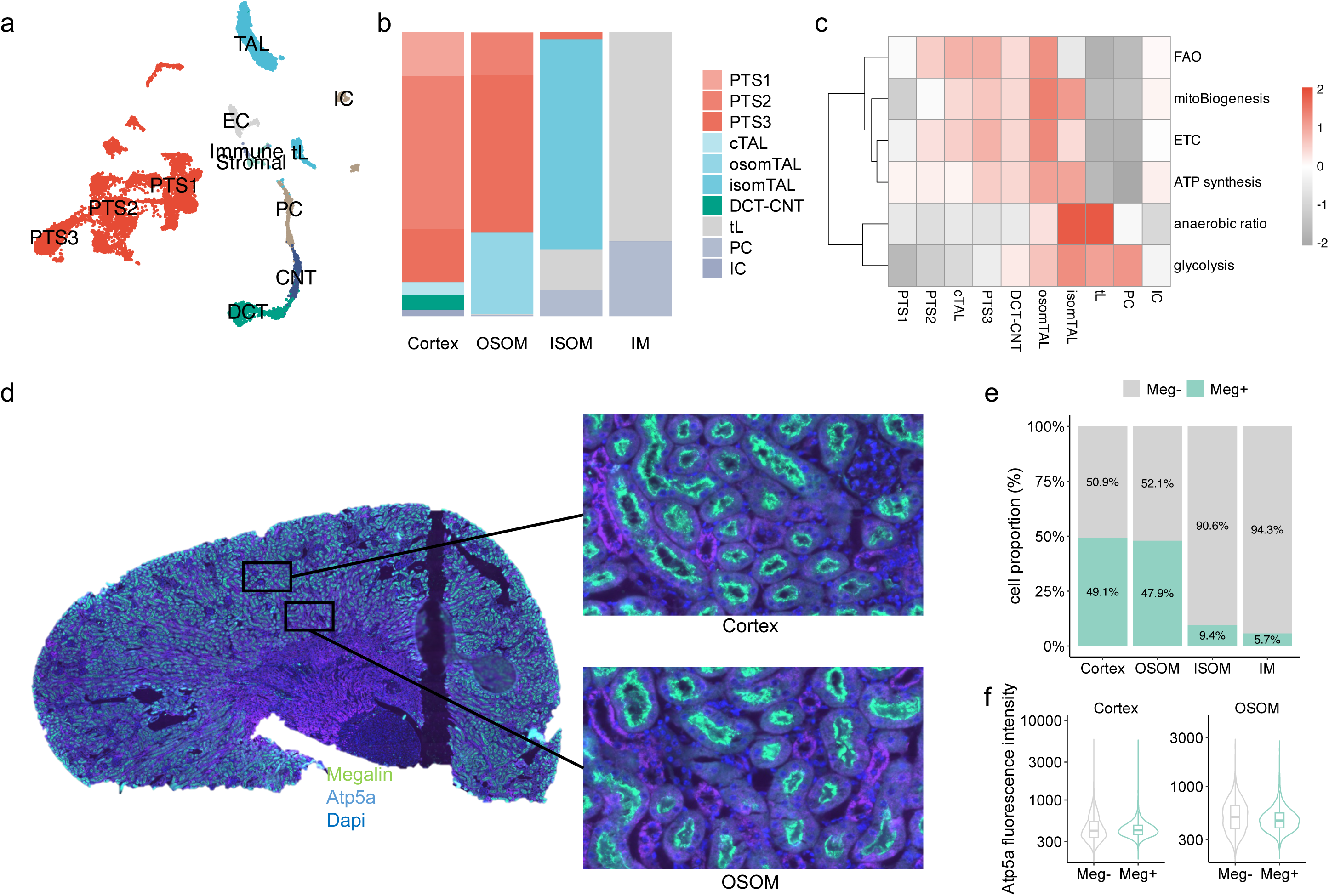
Renal energy metabolism at the cell type level. **a**) UMAP projection of the integrated snRNAseq dataset, showing major renal cell types. **b**) Barplots showing the relative abundance of major renal cell types in each compartment. **c**) Heatmap showing the level of metabolic processes across renal cell types. **d**) Representative immunofluorescence image of Atp5a/megalin co-staining in healthy mice kidney section with a focus on cortex and outer strip of outer medulla (OSOM). **e**) Relative proportion of cells expressing or not megalin across each compartment. f) Violin plots showing the fluorescence intensity level if the Atp5a staining in cells expressing megalin or not in cortex and OSOM. Violin plots display distribution of the data, boxplots display median with interquartile range and dots display mean. FAO fatty acid oxidation; ETC electron transport chain; PS1, PTS2 and PTS3 segment 1, 2 and 3 of the proximal tubules; DTL-ATL, descending and ascending thin limb, TAL thick ascending limb; DCT-CNT, distal convoluted and connecting tubules; EC, endothelial cell; UMAP, uniform manifold approximation and projection.

**Figure 3:**
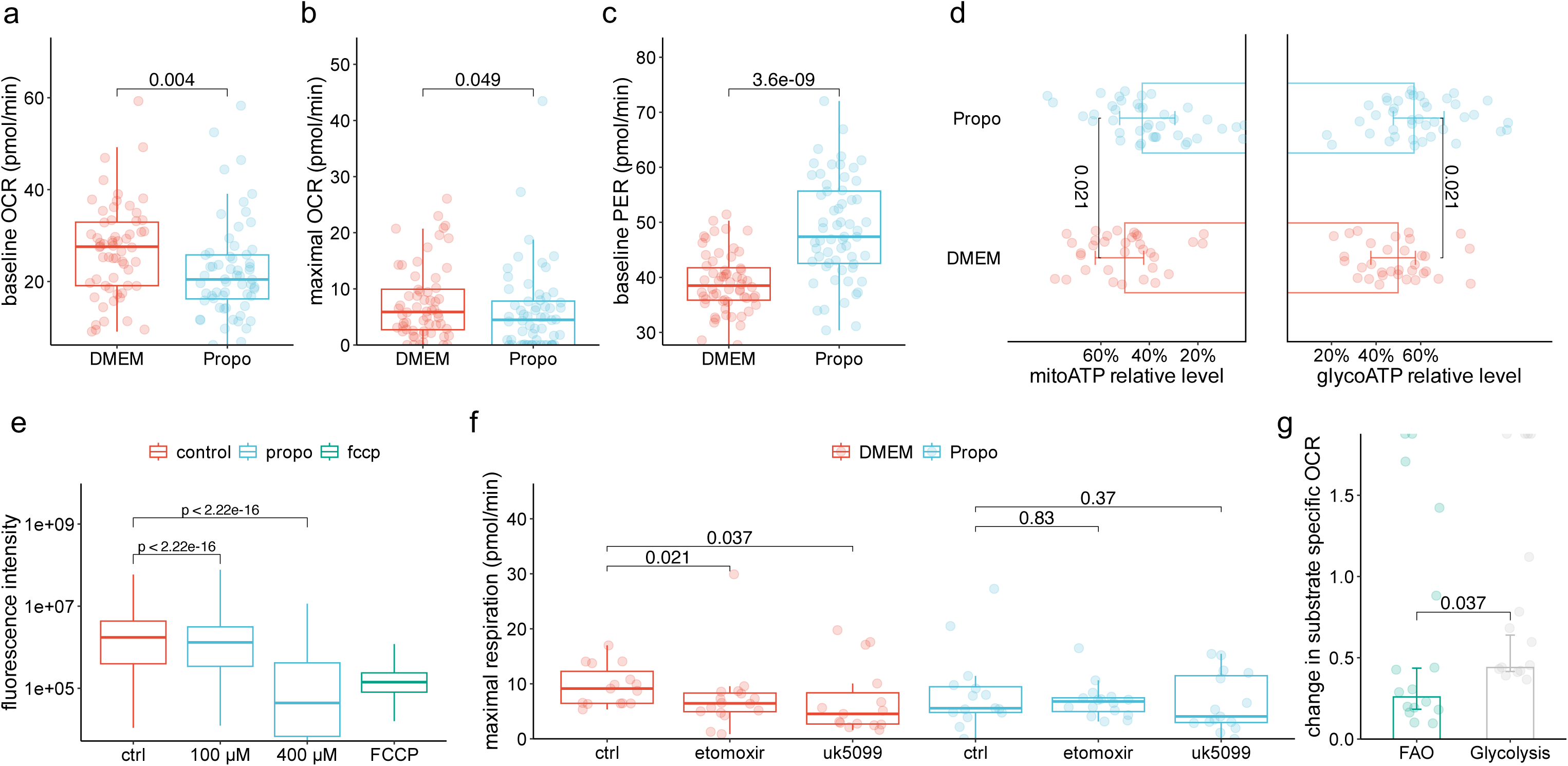
Effect of propofol renal energy metabolism in vitro. **a-c**) Seahorse experiments with (**a**) baseline oxygen consumption rate (OCR in pmol/min), (**b**) maximal OCR (pmol/min) and (**c**) glycolytic proton efflux rate (PER in pmol/min) of primary renal cells treated or not with propofol (Propo). **d**) Barplots showing the relative contribution of oxidative phosphorylation and anaerobic glycolysis to ATP production in renal primary cells treated or not with propofol. **e**) Boxplots showing the mitochondrial membrane potential of HK2 cells treated with different concentrations of propofol or FCCP as positive control, assessed by mitotracker probe and measured using flow cytometry. **f**) Maximal respiration (in pmol/min) of renal primary cells treated or not by propofol with the addition or not of etomoxir to inhibit FAO or uk5099 to inhibit pyruvate metabolism. **g**) Barplots showing the relative change in OCR after propofol treatment related to FAO or aerobic glycolysis decrease. Boxplots and barplots display median with interquartile range.

### Propofol impairs renal mitochondrial function

Given the high rate of oxidative phosphorylation in the OSOM, we wondered whether this area might be particularly sensitive to mitochondrial impairment. We therefore searched for a drug commonly used in clinical practice that has mitochondrial inhibitory properties. Sedatives are one of the three most commonly administered classes of drugs in the ICU setting^42–44^. Of the available sedatives, propofol is the most commonly used agent^45–47^, in line with current guidelines recommending propofol over midazolam to decrease ICU length of stay, duration of mechanical ventilation and delirium^48,49^. Propofol has been shown to penetrate the mitochondrion^50^ and is listed as a drug that interferes with oxidative phosphorylation^51^. Propofol has been shown to affect mitochondria in several organs, including skeletal muscle, heart and liver^52–56^. In the same vein, the most dangerous side effect of propofol is propofol-related infusion syndrome (PRIS), whose pathophysiology involves mitochondrial impairment and decreased fatty acid oxidation^57^. We first investigated whether propofol could also affect mitochondrial function in the kidney. We used the Seahorse platform with primary kidney cells incubated with or without propofol (100 µM). Propofol decreased both baseline and maximal oxygen consumption rate (OCR) and increased proton efflux rate (**Figure 3a-c**). Consistently, we observed a switch in ATP production from oxidative phosphorylation (mitoATP) to anaerobic glycolysis (glycoATP) in propofol-treated kidney cells (**Figure 3d**). We confirmed these findings in the HK2 renal cell line (**Supplemental Figure 4**). These results suggest a decrease in mitochondrial function in renal cells treated by propofol. This was corroborated by a dose-dependent reduction in mitochondrial membrane potential, in HK2 cells treated by propofol-(**Figure 3e**).

To assess whether propofol differentially impairs aerobic glycolysis (*i.e.,* pyruvate metabolism) or fatty acid oxidation (FAO), primary renal cells incubated with or without propofol were treated with etomoxir or UK5099 to inhibit FAO or mitochondrial pyruvate metabolism, respectively. In primary renal cells, both etomoxir and uk5099 reduced OCR, suggesting that these cells rely on both FAO and aerobic glycolysis to meet ATP demand. In contrast, neither etomoxir nor uk5099 decreased OCR in propofol-treated primary kidney cells in line with an inhibition of oxidative phosphorylation (**Figure 3f**). Combining these results, we showed that propofol primarily impairs FAO (**Figure 3g**).

To extend this finding to the whole kidney, we used an *ex vivo* kidney perfusion platform previously described (**Figure 4a**)^34^ and firstly confirmed the absence of drift over time in baseline condition (**Supplemental Figure 5**). After 15 minutes of equilibration, we added propofol (100 µM) to the perfusion solution. We found a decrease in renal oxygen consumption in the kidney after propofol infusion, in line with a decrease in the mitochondrial function (**Figure 4b**). Furthermore, we observed an increase in renal glucose uptake and lactate release, which collectively indicate a compensatory glycolytic switch (**Figure 4c, d**). Additionally, the ratio between carbon dioxide and oxygen arteriovenous difference increased after propofol infusion, suggesting a change in substrates from fatty acids to glucose (**Figure 4e**).

**Figure 4:**
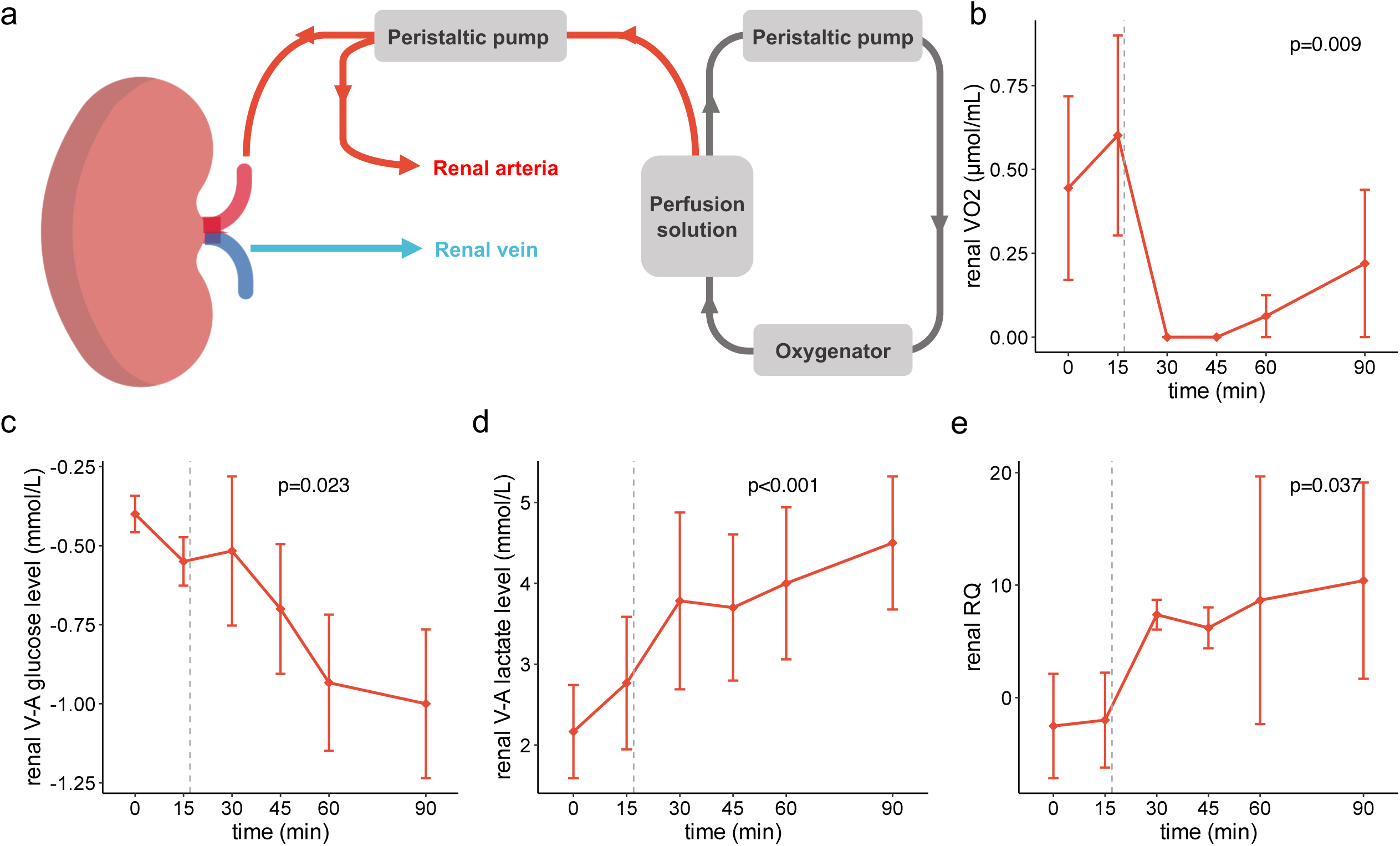
Effect of propofol on renal energy metabolism ex vivo. **a**) Schematic representation of the ex vivo kidney perfusion platform. **b**) Renal oxygen consumption (in µmol/min) before and after propofol infusion at 15 minutes. **c, d**) Renal arteriovenous difference in glucose (mmol/L) (**c**) and lactate (mmol/L) (**d**) before and after propofol infusion at 15 minutes. **e)** Renal respiratory quotient (RQ) before and after propofol infusion at 15 minutes. Error bars display mean ± standard error.

Finally, we observed that peritoneal injection of propofol (150 mg/kg) resulted in elevated lactate levels in healthy mice compared to midazolam (20 mg/kg) (**Supplemental Figure 6**).

Collectively, these findings indicate that propofol impairs mitochondrial function and oxidative pathways including FAO and pyruvate metabolism in the kidneys.

### Preconditioning with propofol worsens tubular injury during ischemia reperfusion

To investigate whether a reduction in mitochondrial respiration could impact the course of renal injury and early recovery, we performed a renal unilateral ischaemia reperfusion injury (IRI) in mice that had been preconditioned either by propofol (150 mg/kg) or midazolam (20 mg/kg), the second most commonly used sedative in intensive care, with no effect on oxidative metabolism in vitro (**Supplemental Figure 4**). To avoid a bias due to sedation ^35^, all mice received isoflurane from preconditioning until the end of ischaemia (**Figure 5a**). Two days after the injury, there was no difference in renal function, assessed by measured glomerular filtration rate (GFR), serum urea and serum creatinine levels (**Supplemental Figure 7**). However, we observed higher tubular injury on kidney slices, assessed by digital tissue detection script. Interestingly, only the OSOM display significant differences (**Figure 5b, c, Supplemental Figure 8**). Similarly, we found a decreased expression of TFAM, a marker of mitochondrial biogenesis (**Figure 5d,e**) and a higher expression of HK1, a glycolytic enzyme (**Figure 5f,g**) in the OSOM of mice preconditioned with propofol. Interestingly, when distinguishing cells that either did or did not express megalin, the decrease in TFAM expression was significant only in cells that did not express megalin, suggesting that propofol preferentially impairs non-proximal tubule segments (**Supplementary Figure 9**).

**Figure 5:**
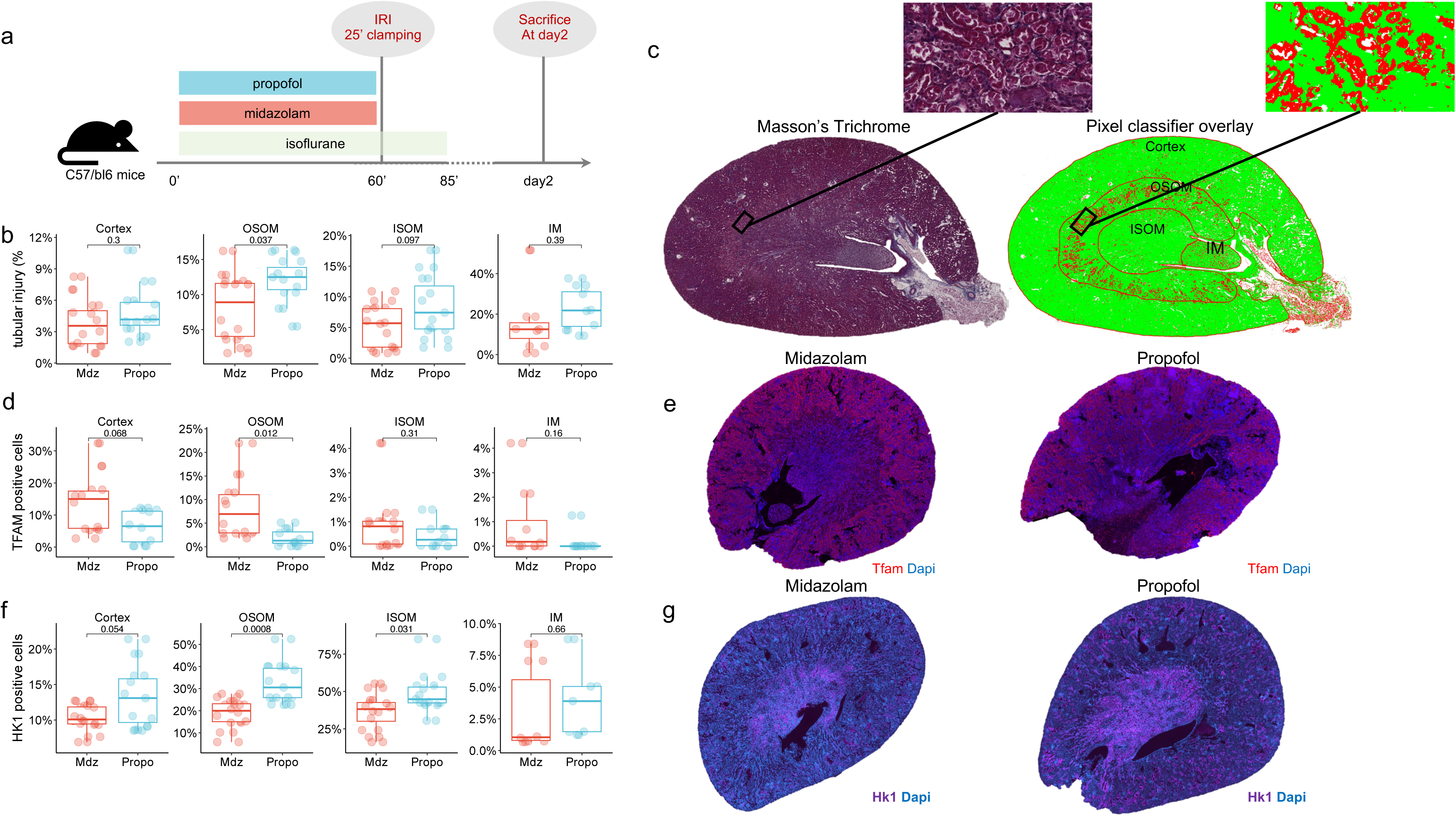
Effect of propofol preconditioning on acute kidney injury. **a**) Schematic representation of the experimental protocol. **b**) Quantification of the tubular injury in the cortex, OSOM, ISOM and IM across animals preconditioned by midazolam or propofol two days after ischemia-reperfusion injury **c**) Representative Masson’s trichrome staining showing tubular fibrosis (left panel) with overlaid pixel classifier detecting tubular injury (right panel)) **d)** Quantification of the TFAM positive cells in the cortex, OSOM, ISOM) and IM across animals preconditioned by midazolam or propofol two days after ischemia-reperfusion injury. **e**) Representative immunofluorescence images of TFAM staining in kidney preconditioned by midazolam or propofol and quantification of the TFAM positive cells in the cortex, OSOM, ISOM and inner medulla across animals preconditioned by midazolam or propofol two days after ischemia-reperfusion injury. **f**) Quantification of the HK1 positive cells in the cortex, OSOM, ISOM) and IM across animals preconditioned by midazolam or propofol two days after ischemia-reperfusion injury. **g**) Representative immunofluorescence images of HK1 staining in kidney preconditioned by midazolam or propofol and quantification of the TFAM positive cells in the cortex, OSOM, ISOM and inner medulla across animals preconditioned by midazolam or propofol two days after ischemia-reperfusion injury. Boxplots display median with interquartile range. C Cortex; OSOM Outer strip outer medulla; ISOM Inner strip outer medulla; IM Inner medulla

These results suggest that the high oxidative phosphorylation activity of the OSOM makes it vulnerable to mitochondrial impairment upon ischemic injury, leading to a metabolic switch toward anaerobic glycolysis and exacerbated tubular injury. By targeting FAO inhibition, propofol may also worsen tubular adaptability to metabolic disturbances and response to acute injury.

### Propofol Impairs Mitochondrial Respiration in the Human Kidney

To apply these findings to patients, we took advantage of our cohort, which included 42 kidney allograft recipients. Each of these patients underwent protocol renal biopsies before implantation, after reperfusion and at 3 and 12 months. For each sample and each time, we assessed energy metabolism using transcriptomic data. These data were further evaluated for association with the cumulative dose of propofol received during transplantation. Our findings indicate that propofol is associated with decreased oxidative metabolism, including FAO, TCA, and ETC between biopsies sampled before and immediately after transplantation (**Figure 6a**). Additionally, we explored the association between the cumulative dose of propofol and renal allograft outcomes. Patients exhibiting a Banff Lesion ct score of greater than 1 at one year had received a higher dose of propofol during transplantation (**Figure 6b**). In a similar vein, propofol was associated with worse renal function at one year and with a higher loss of renal function between three and 12 months (**Figure 6c, d**).

**Figure 6.**
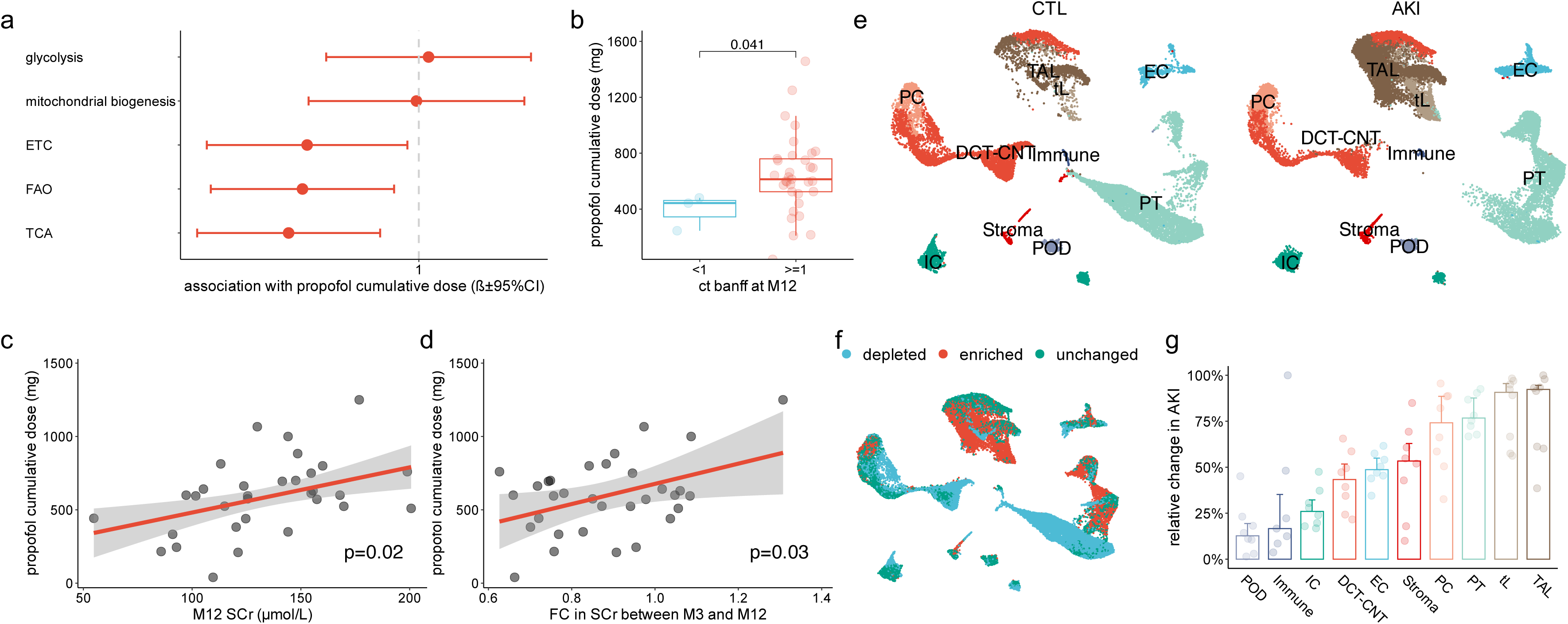
Propofol in human kidney allograft recipients and change in cell subpopulations during AKI. **a**) forest plot showing the association (ß coefficient with 95% confidence interval) between cumulative dose of propofol (mg) received during transplantation and change in renal energy metabolism assessed with singscore during transplantation. **b**) cumulative dose of propofol (mg) received during transplantation among patients displaying a Banff ct score higher or lower than 1 at one year. **c, d**) scatter plot showing the association between cumulative dose of propofol (mg) received during transplantation and serum creatinine level (µmol/L) at one year (**c**) or change in serum creatinine levels between 3 months and one year after transplantation (**d**). (**e-f**) UMAP projection of the integrated snRNAseq human split by AKI status (**e**) and showing the subpopulation of cells enriched, depleted or unchanged between AKI and control (**f**). (**g**) barplots showing the relative change in subpopulation across cell types in AKI patients. Immune, immune cells; POD, podocytes; IC, intercalated cells; CNT, connecting tubule cells; DCT, distal convoluted tubule cells; PT, proximal tubule cells; EC, endothelial cells; Stroma, stromal cells; PC, principal cells; LOH, loop of Henle cells. Barplots display median with interquartile range.

Taken together, we found an association between the cumulative dose of propofol received during transplantation and decreased oxidative metabolism and the worst renal outcomes at one year in renal allograft recipients.

### Not only PT cells but TAL cells are also significantly targeted by injury in AKI

To ascertain whether TAL, the predominant segment of the OSOM, is particularly susceptible to injury during AKI, we obtained kidney biopsies from five ICU patients experiencing AKI in the context of multi-organ failure before planned withdrawal of resuscitation measures, as previously described^40^. Major cell types were identified and evident changes in subpopulations was observed among AKI and Control patients (**Figure 6e**). DA-seq was used to detect cell subpopulations with differential abundance between AKI and control patients (**Figure 6f**). Among the segments characterized, the loop of Henle exhibited the greatest degree of relative change (depletion or enrichment) during AKI (**Figure 6g**). Similar results were found in the largest human snRNAseq dataset^58^ (**Supplemental Figure 10**), in accordance with previous results^59^.

## Discussion

The kidneys are highly sophisticated organs showing a complex architecture with different cell types in distinct locations. Each of the 4 main regions (histological layers) of the kidney has a specific pattern of cell composition and metabolism (oxygen consumption, and substrate preference) according to its specific function and microenvironment (oxygen supply). While renal mitochondrial and FAO dysfunction have emerged as key players in the pathophysiology of AKI^13–21^, it is unclear whether metabolic activity could be linked to tubular injury severity in a regional manner. This study highlighted the high energy metabolic rate of the OSOM, mainly PTS3, TAL and DCT-CNT. These cell types rely on both aerobic and anaerobic processes, including FAO and glycolysis to meet ATP demands. Therefore, we reported that propofol, by impairing mitochondrial oxidative phosphorylation and FAO, worsens tubular injury in this area during acute kidney injury.

The bulk of ATP synthesized in the kidney derives from oxidative phosphorylation. However, the spatial heterogeneity of renal energy metabolism in terms of rate of activity and preferred substrates is well documented and is related to the different functions of each nephron segment^23,28,60,61^. There is a strong link between active transport and mitochondrial respiration^62,63^, with inhibition of sodium reabsorption reducing renal oxygen consumption^64,65^. This reflects the coupling between ATP production and consumption^66^. Consequently, the distribution of sodium pumps along the nephron is not uniform, with the highest density observed in the TAL, followed by the DCT and proximal tubules^23,60,66,67^. Similarly, the enzymes involved in energy production show maximal activity in the TAL and DCT ^23,24,68^. The expression of TCA cycle enzymes^23,27,69^ and the mitochondrial density^26,27,69^ are also maximal in these segments, exceeding that observed in the proximal tubules. At the functional level, oxidation rate, as measured by labeled CO₂ production after incubation with U-14C substrates, has also been reported to be maximal in the DCT, TAL and proximal tubule ^70^. In this study, using transcriptomics approaches, we demonstrated that the PTS3, TAL and DCT-CNT dwelling within the OSOM are the most metabolically active parts of the nephron.

In addition, the substrates used to make ATP differ throughout the nephron. FAO has been shown to be particularly intense in the OSOM. At the protein expression level, carnitine palmitoyltransferases, a key FAO enzyme, is particularly well expressed in the TAL and DCT^71^. The expression ratio between ß-hydroxyacyl-CoA dehydrogenase and citrate synthase is maximal in the late proximal tubule^69^. Functionally, the OSOM oxidizes palmitate at a higher rate than the cortex and ISOM^72,73^. ATP production from hydroxybutyrate, a ketone produced during FAO occurs in the PTS3, TAL and DCT but not in the S1 and S2 parts of the proximal tubule ^11^. Recently, a group coupled stable isotope-labelled nutrient infusion to matrix-assisted laser desorption ionization imaging mass spectrometry to quantify metabolic activity in the mouse kidney. Lactate and free fatty acids were found to be the major direct contributors to the medullary TCA cycle ^74^. In contrast, glycolysis is absent in the early proximal tubules (pars convolute) but appears later in the pars recta and progressively increases until the distal nephron ^11,69,70,73,75–77^. It is noteworthy that glucose undergoes different catabolic pathways. While it is mostly oxidized in the PTS3, TAL and DCT segments, it is mostly anaerobically converted to lactate in the ISOM segments and in the inner medulla ^11,69,70,73,75,76^. This is consistent with our findings, which showed a high FAO rate in the late part of PTS3, the DCT-CNT and the cortical and OSOM TAL. We also found that glycolysis starts in the TAL, is initially aerobic, and becomes anaerobic in the inner medulla.

Renal mitochondrial function inhibition induced by propofol was expected, as a decrease in mitochondrial membrane potential and mitochondrial oxygen consumption after incubation with propofol has been consistently described in other organs ^52–56^. Propofol has been shown to reduce the activity of ETC enzymes due to its structural similarity to ubiquinone and the cytochrome aa3 subunit, which allows propofol to accept but not to transfer electrons. This electron transfer interference results in disruption of the ETC and prevention of mitochondrial membrane potential formation ^54,78–81^. At higher concentrations, propofol has also been reported to affect all mitochondrial ETC complexes (*i.e.,* I, II, III and IV)^52,53,79,82^. Consistently with these findings, propofol reduced oxygen consumption in a perfused guinea pig heart model by decreasing electron flux along the respiratory chain^83^. In addition to these effects on ETC, propofol may also have a direct effect on fatty acid oxidation. This hypothesis was first proposed in 2001 in a two-year-old child with propofol infusion syndrome who had elevated serum levels of malonyl carnitine and C5 acyl carnitine, two early metabolites of FAO^84^. Recently, another group reported a decrease in FAO rate, assessed by [1-^14^C]palmitate isotope assay^53^. The FAO defect was seen at extremely low concentrations of propofol, before reaching the levels associated with ETC reduction. At the enzymatic level, propofol has been shown to interfere with FAO between CPT-2 and the entry of electrons from FAO to ETC via complex II ^53,83^. Surprisingly, some groups have reported a nephroprotective effect of propofol ^85–89^. Three factors may explain this discrepancy. First, the intraperitoneal route of propofol administration resulted in a longer infusion time compared to the intravenous route ^90^. Secondly, propofol was administered 60 min before IRI, resulting in a longer preconditioning period compared to the other studies. Thirdly, and most importantly, all earlier studies compared propofol with saline. Our recent findings have shown that anaesthesia confers renal protective effects related to the modification of the IRI transcriptomic profile ^35^. In this context, the reported propofol-related renal protection may be due to sedation and not to propofol itself. In our study, we used midazolam as comparison and administered isoflurane to all mice to ensure an equivalent level of sedation.

Why propofol mainly affects the OSOM is of great interest. We hypothesize that the high metabolic demand of the OSOM makes it very sensitive to a mild inhibition of mitochondrial function. Similarly, by impairing FAO, propofol may have specifically affected the OSOM, where FAO occurs at a high rate. Furthermore, the sensitivity to mitochondrial inhibition does not seem to be uniform along the nephron. The PTS3 and TAL segments have been shown to have a more pronounced reduction in ATP production following mitochondrial inhibition by antimycin-A than other segments, despite increased compensatory glycolysis ^91^. The heterogeneity of renal oxygenation, which has been well documented in the literature ^92^, may also have exacerbated the vulnerability of the OSOM. Finally, the efficiency of oxygen utilization for oxidative phosphorylation is lower in the TAL than in the proximal tubule ^93^, meaning that they require more oxygen to synthesize ATP, increasing its vulnerability.

In conclusion, we demonstrated that the OSOM is the most metabolically active and vulnerable region of the kidney upon acute injury using spatial multimodal transcriptomics in mice and humans. Moreover, as shown by extrarenal studies, we proved that propofol impairs oxidative phosphorylation and FAO in the kidney by targeting the OSOM (composed mainly of the S3 segment of the proximal tubule and the thick ascending limb of the loop of Henle), which depend on oxidative phosphorylation for ATP production and have high energy demands. Although not detected at the GFR level, the detrimental effect of propofol on metabolism and injury severity may impair renal recovery and promote further progression to chronic kidney disease.

## Disclosure

The authors declare that they have no competing interests.

## Funding

DL is supported by two young researcher grants from the Geneva University Hospitals (PRD 5-2020-I and PRD 4-2021-II).

## Acknowledgements

We thank the BioMedTech Facilities, INSERM US36, CNRS UAR2009, Université Paris Cite for its help.

## Authors’ contributions

Conceptualization, D.L., C.V.; methodology, D.L., P.G., A.G.; formal analysis, D.L., A.G., J.S., M.T..; experiment, T.V., J.S., F.S., M.F., S.P.; writing—original draft preparation, D.L.; writing—review and editing, A.G., T.V., J.S., M.T., F.S., SdS, E.F., SNK, P.G., D.L.; supervision, D.L. All authors have read and agreed to the published version of the manuscript.

## Data Sharing Statement

The datasets used and or analyzed during the current study are available from the corresponding author on reasonable request. The scRNAseq dataset was downloaded from GEO at the accession number GSE131882.

---

**Supplemental Figure 1:**
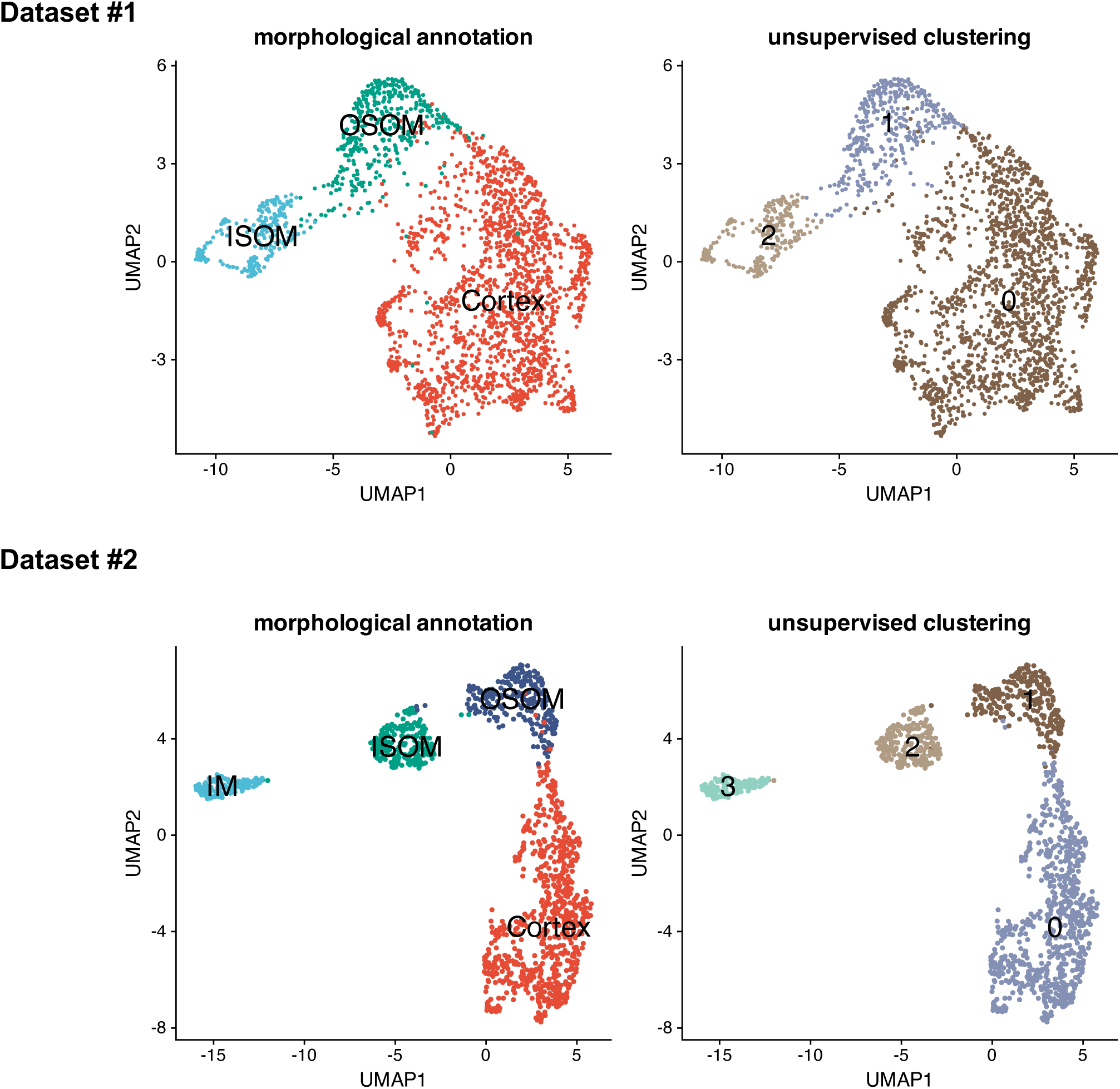
Unsupervised clustering overlaps with morphological annotation: UMAP projection of the integrated snRNAseq dataset and annotated using unsupervised clustering (left panel) or morphological based annotation (right panel) in the two spatial transcriptomic datasets.

**Supplemental Figure 2:**
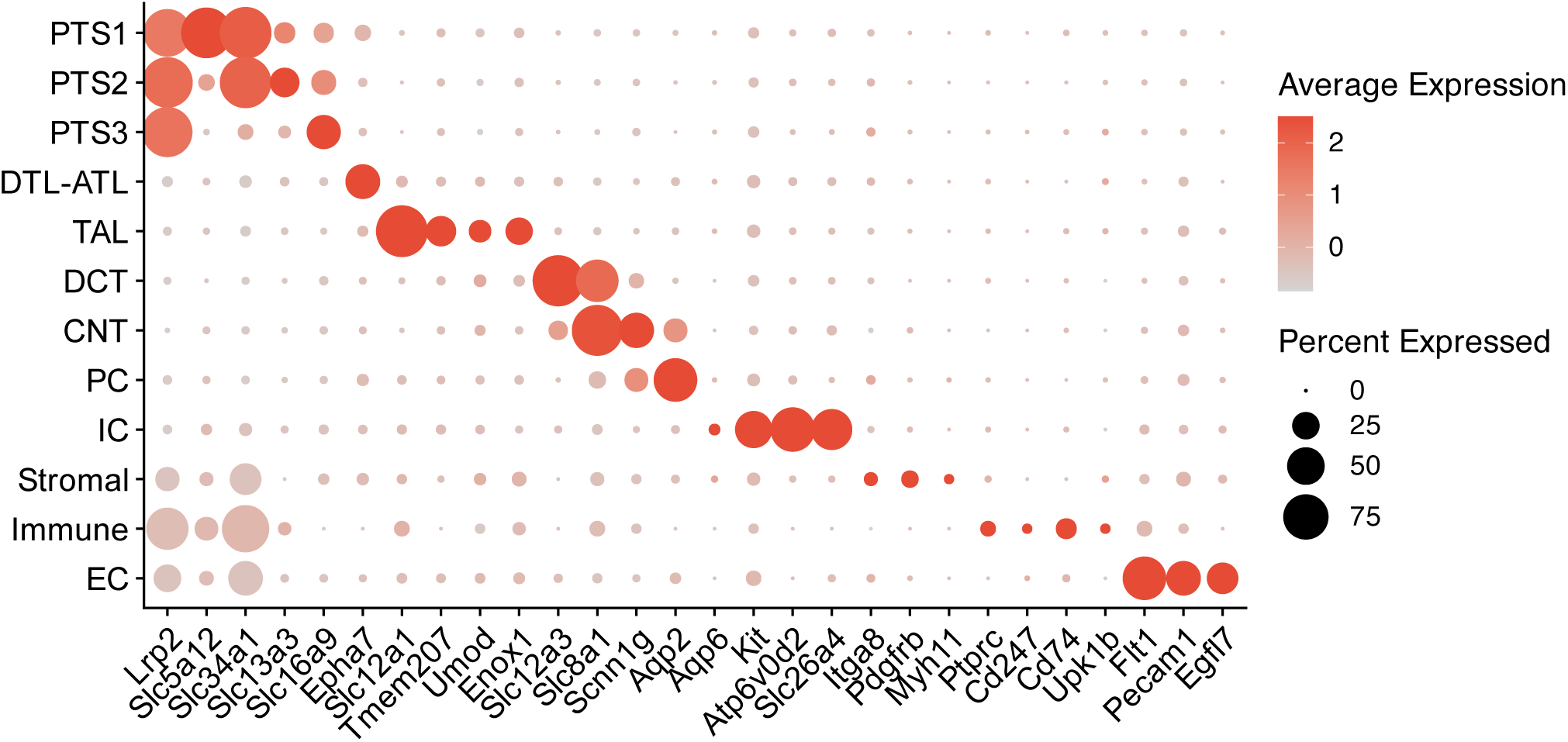
Identification of major renal cell types: Dot plot showing mean marker gene expression values and proportion expressed for each renal cell type.

**Supplemental Figure 3:**
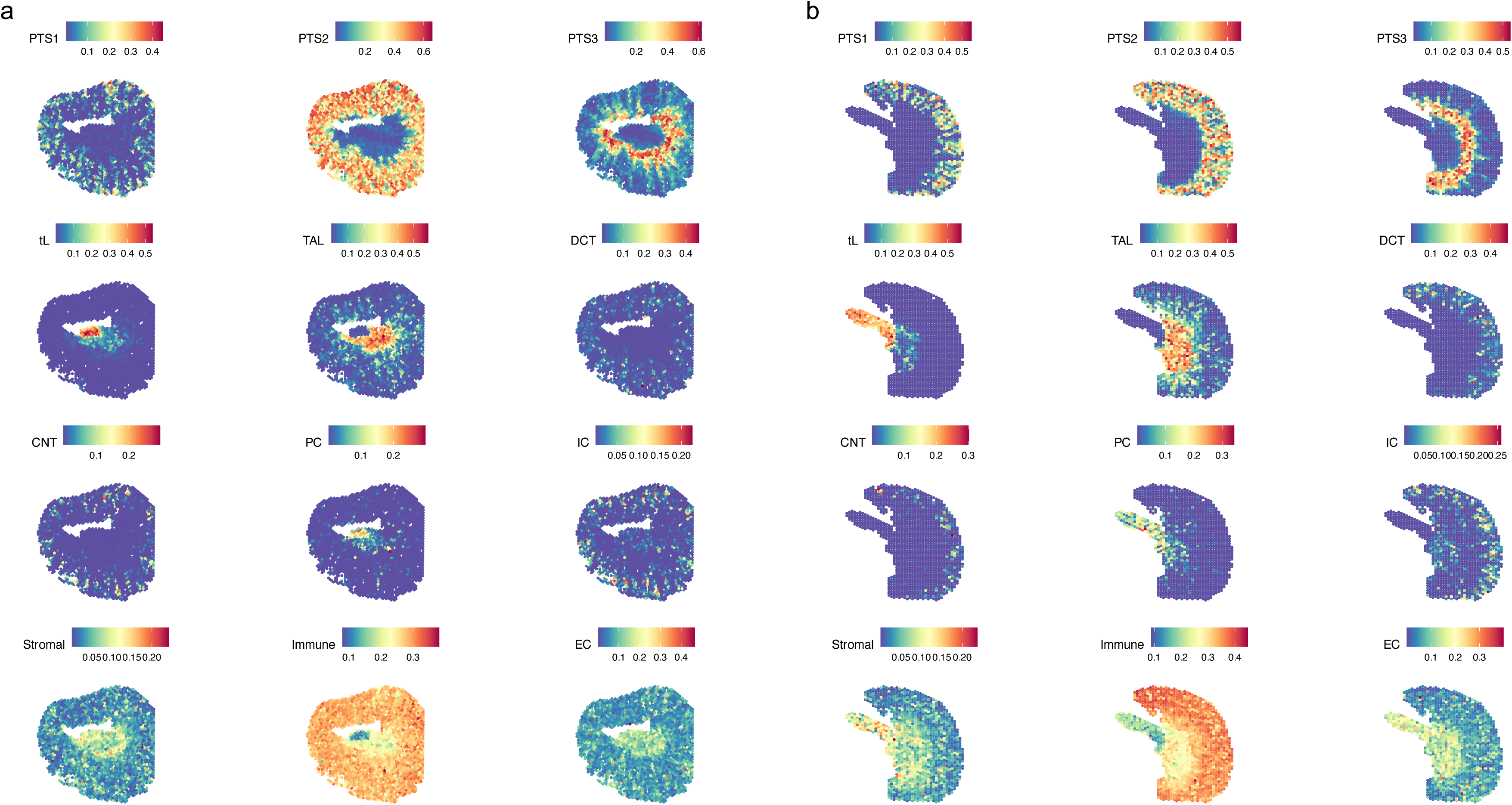
Cell type decomposition: Spatial feature plot showing the cell type assigned by robust cell decomposition for each spot in the datasets #1 (**a**) and #2 (**b**).

**Supplemental Figure 4:**
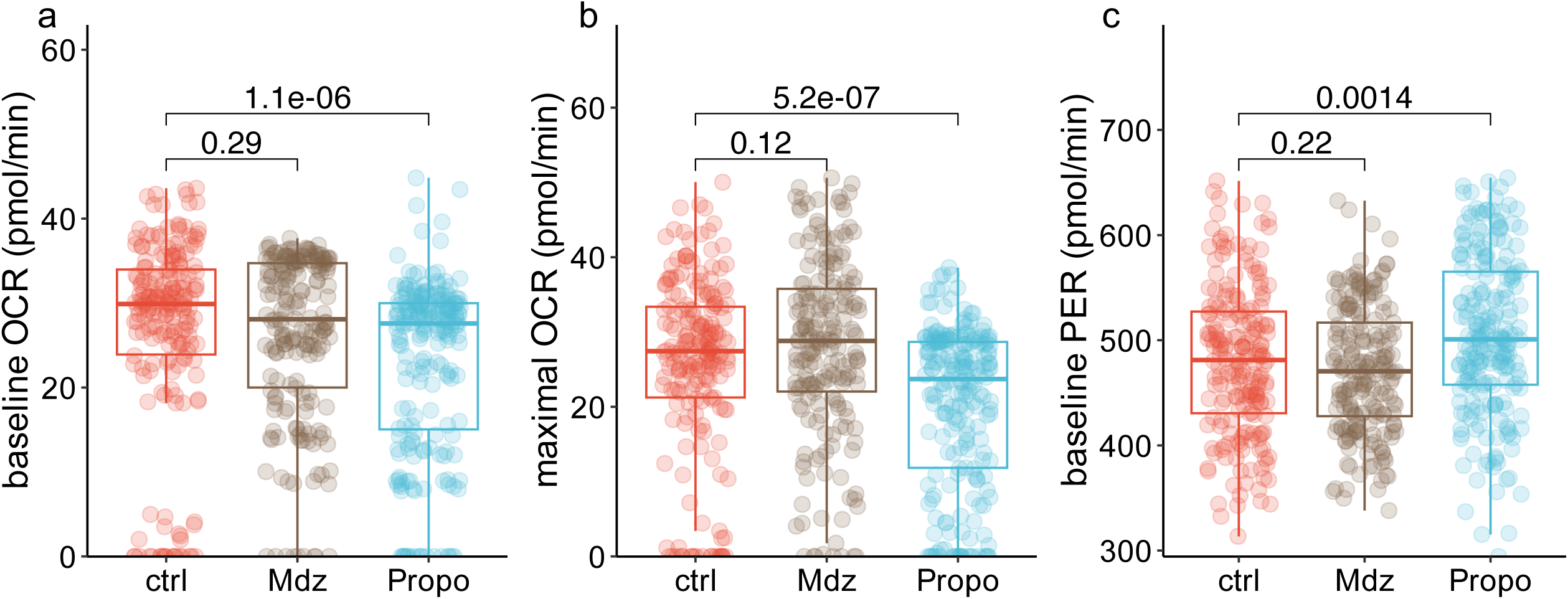
Effect of propofol on renal energy metabolism in HK2 cells. **a**) baseline oxygen consumption rate (OCR in pmol/min), (**b**) maximal OCR (pmol/min) and (**c**) glycolytic proton efflux rate (PER in pmol/min) of HK2 cells untreated or treated with midazolam (1 µM) or propofol (100 µM). Boxplots display median with interquartile range.

**Supplemental Figure 5:**
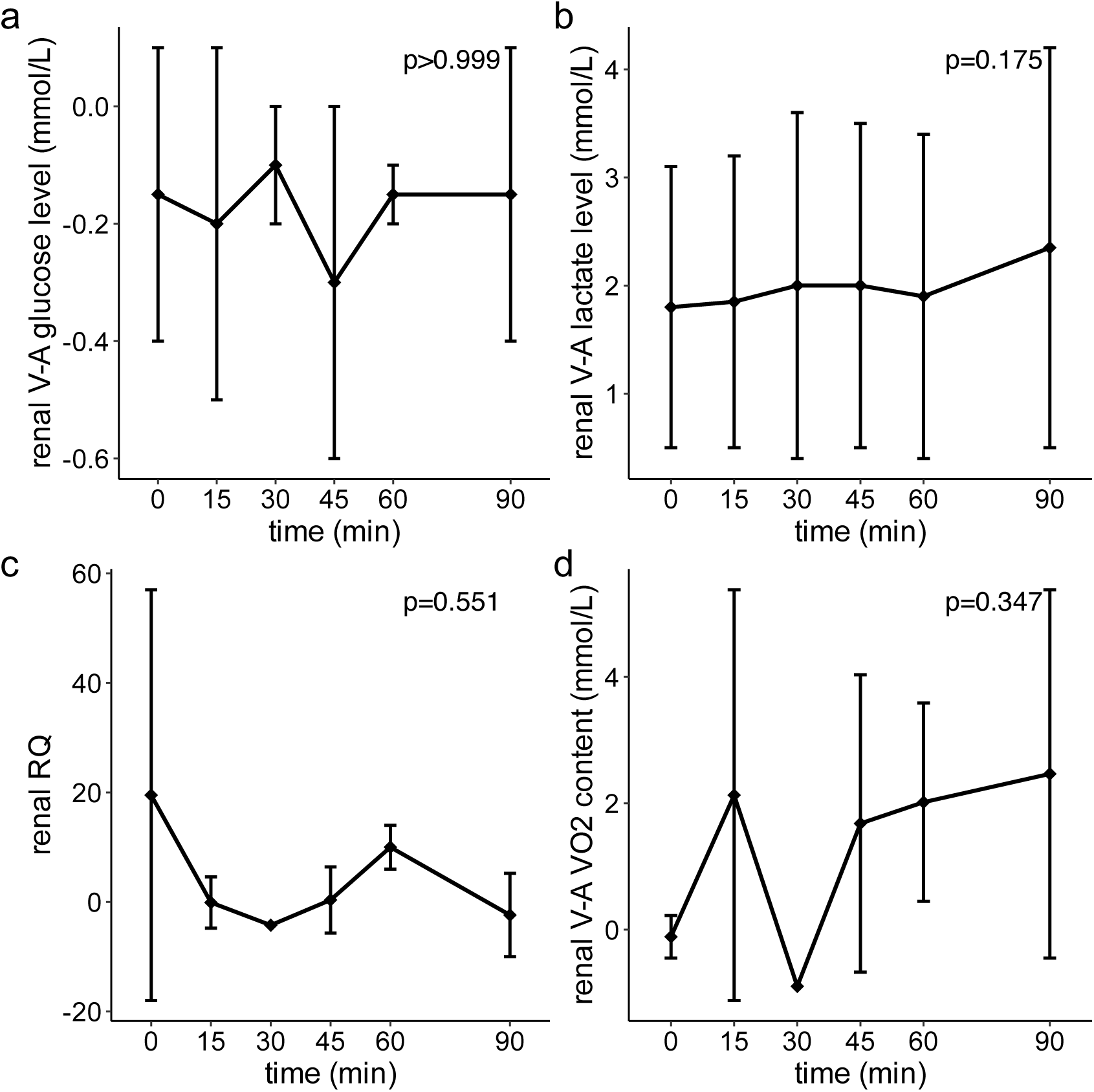
Time control for ex vivo renal perfusion. Renal arteriovenous difference in glucose (**a**) and lactate (**b**), renal respiratory quotient (**c**) and renal oxygen consumption over time during the ex vivo renal perfusion assay. Error bars display mean ± standard error.

**Supplemental Figure 6:**
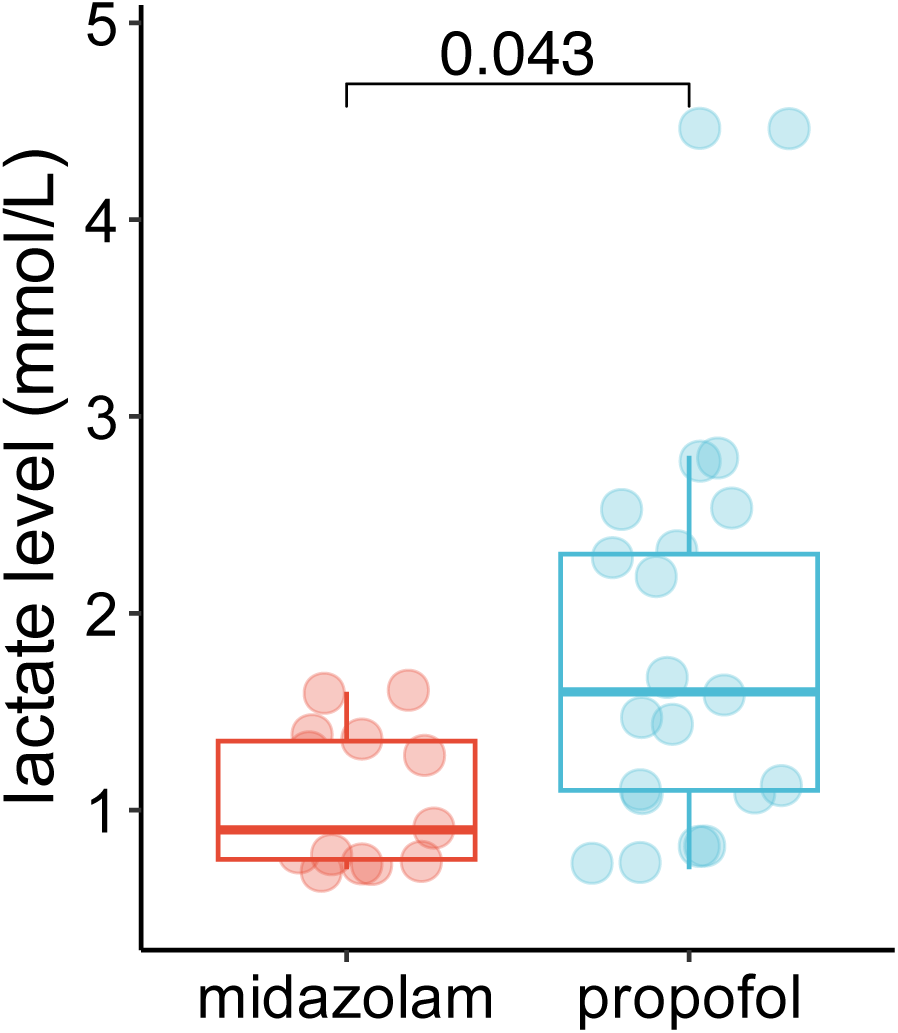
Capillary lactate levels after intraperitoneal injection of midazolam or propofol. Boxplots display median with interquartile range.

**Supplemental Figure 7:**
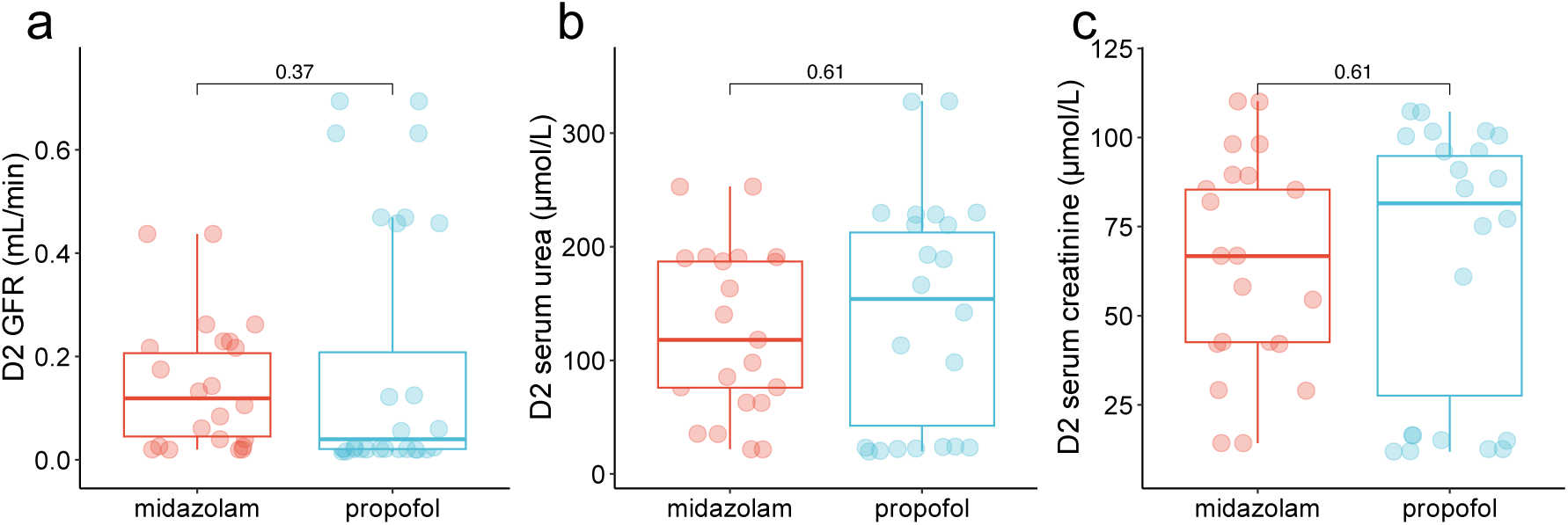
Renal function on day 2 after IRI. Measured glomerular filtration rate (GFR in mL/min) (**a**), serum urea (µmol/L) (**b**) and creatinine levels (µmol/L) (**c**) in mice on day 2 after IRI and preconditioned by midazolam or propofol. Boxplots display median with interquartile range.

**Supplemental Figure 8:**
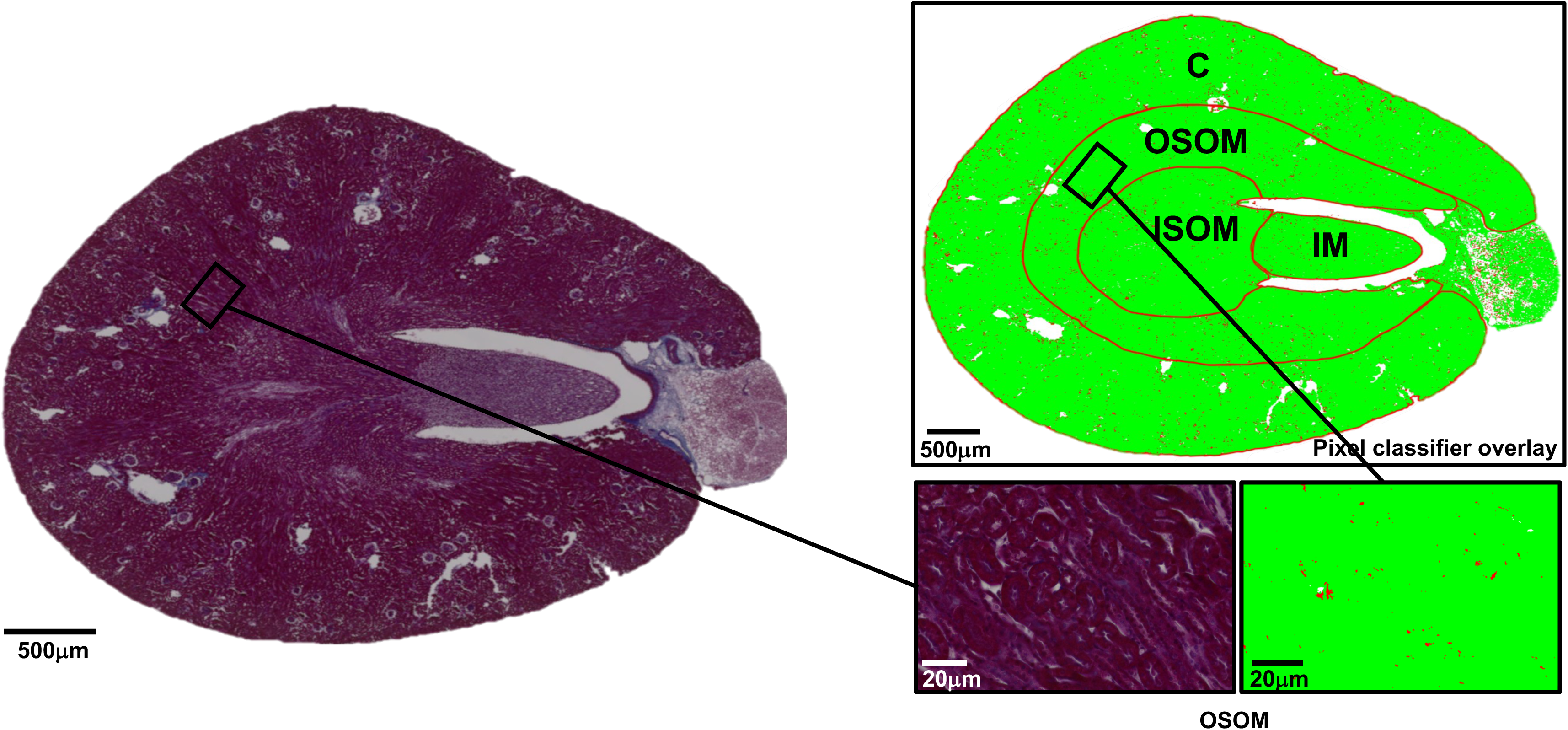
Representative Masson’s trichrome staining with overlaid pixel classifier detecting tubular injury in healthy kidney tissue. C Cortex; OSOM Outer strip outer medulla; ISOM Inner strip outer medulla; IM Inner medulla.

**Supplemental Figure 9:**
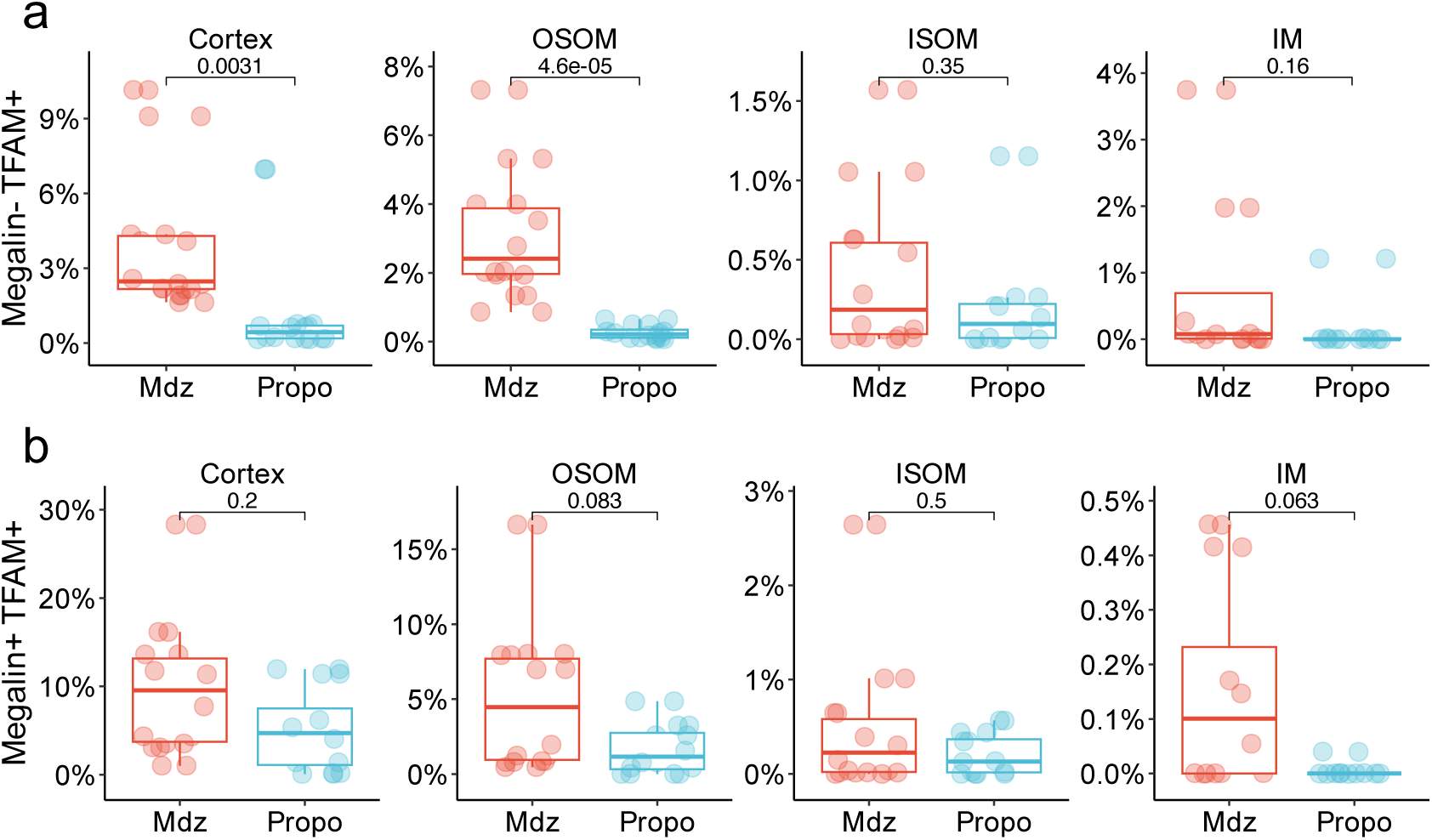
TFAM staining in kidney sections. Relative quantification of cells co-expressing TFAM but not megalin (**a**) or TFAM plus megalin (**b**) in each compartment and in mice two days after IRI and preconditioned with midazolam or propofol. Boxplots display median with interquartile range.

**Supplemental Figure 10:**
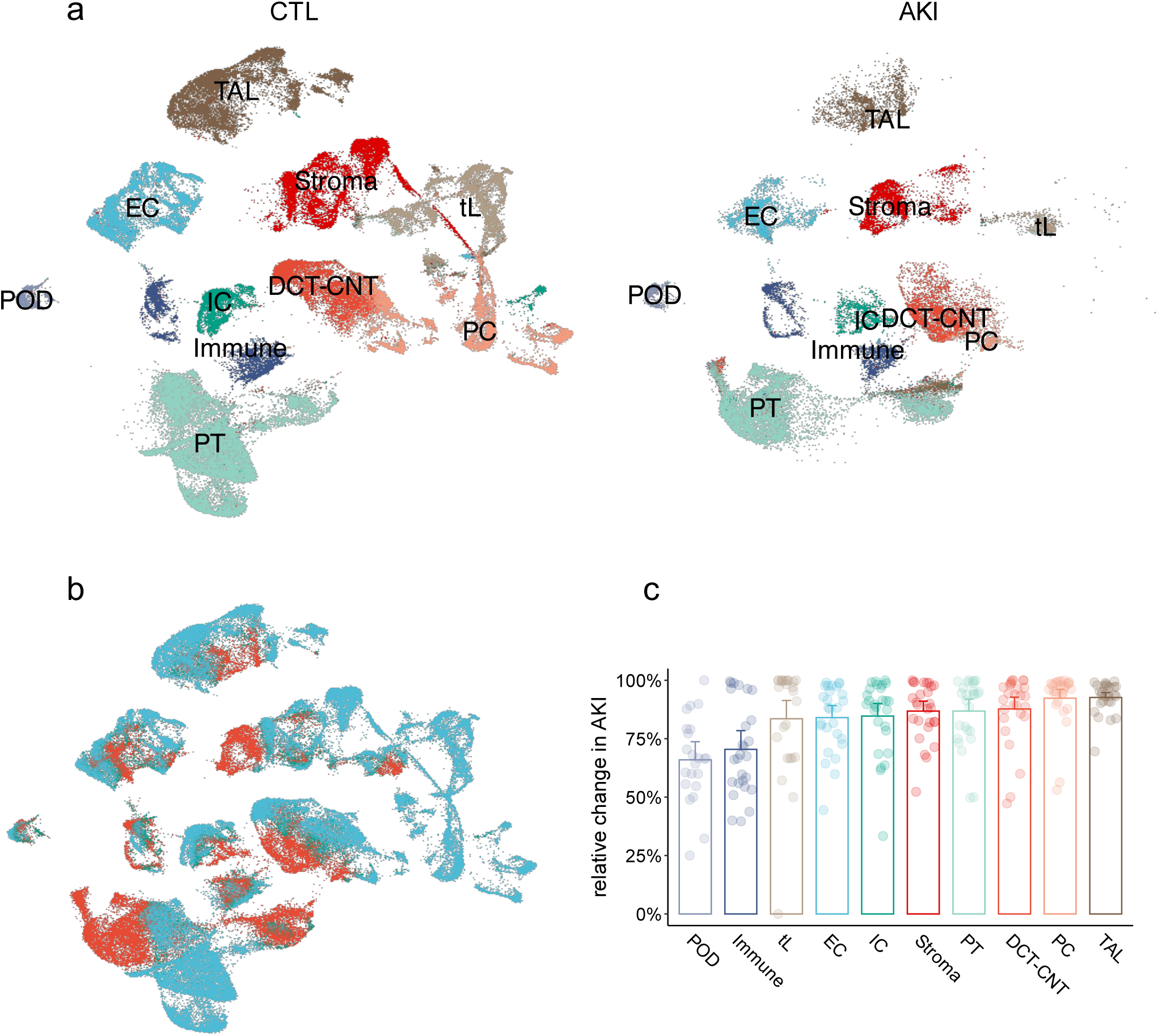
Injured segment in human KPMP dataset. (**a, b**) UMAP projection of the integrated snRNAseq KPMP human split by AKI status (**a**) and showing the subpopulation of cells enriched, depleted or unchanged between AKI and control (**b**). (**c**) Barplots showing the relative change in subpopulation across cell types in AKI patients. Barplots display mean with 95% confidence interval. Immune, immune cells; POD, podocytes; IC, intercalated cells; CNT, connecting tubule cells; DCT, distal convoluted tubule cells; PT, proximal tubule cells; EC, endothelial cells; Stroma, stromal cells; PC, principal cells; LOH, loop of Henle cells.

## Notes

### Competing Interest Statement

The authors have declared no competing interest.

